# Activating and inhibiting nucleotide signals coordinate bacterial anti-phage defense

**DOI:** 10.1101/2025.07.09.663793

**Authors:** Sonomi Yamaguchi, Samantha G. Fernandez, Douglas R. Wassarman, Marlen Lüders, Frank Schwede, Philip J. Kranzusch

## Abstract

The cellular nucleotide pool is a major focal point of the host immune response to viral infection. Immune effector proteins that disrupt the nucleotide pool allow animal and bacterial cells to broadly restrict diverse viruses, but reduced nucleotide availability induces cellular toxicity and can limit host fitness(Ahmad et al., 1998; Goldstone et al., 2011; Hsueh et al., 2022; Itsko & Schaaper, 2014; Tal et al., 2022). Here we discover a bacterial anti-phage defense system named Clover that overcomes this tradeoff by encoding a deoxynucleoside triphosphohydrolase enzyme (CloA) that dynamically responds to both an activating phage cue and an inhibitory nucleotide immune signal produced by a partnering regulatory enzyme (CloB). Analysis of Clover phage restriction in cells and reconstitution of enzymatic function in vitro demonstrate that CloA is a dGTPase that responds to viral enzymes that increase cellular levels of dTTP. To restrain CloA activation in the absence of infection, we show that CloB synthesizes a dTTP-related inhibitory nucleotide signal p3diT (5′-triphosphothymidyl-3′5′-thymidine) that binds to CloA and suppresses activation. Cryo-EM structures of CloA in activated and suppressed states reveal how dTTP and p3diT control distinct allosteric sites and regulate effector function. Our results define how nucleotide signals coordinate both activation and inhibition of antiviral immunity and explain how cells balance defense and immune-mediated toxicity.

## MAIN TEXT

Deoxynucleoside triphosphohydrolase (dNTPase) enzymes are effector proteins in animal and bacterial cells that hydrolyze dNTP nucleotides and inhibit viral replication(Goldstone et al., 2011; Tal et al., 2022). Bacterial anti-phage defense operons are known to encode dNTPase enzymes that restrict a broad range of phages, but these enzymes function through unknown mechanisms of regulation(Tal et al., 2022; Tesson et al., 2022). A recent bioinformatic analysis reported a two-gene system in *Salmonella enterica* SA20044414 that encodes a predicted dNTPase and an accessory nucleotidyltransferase (NTase) enzyme, suggesting the possible existence of cellular signaling dedicated to regulating dNTPase function(Nicastro et al., 2023). We analyzed bacterial anti-phage defense islands and identified >650 related dNTPase-NTase systems across diverse proteobacteria, *Bacillus*, *Clostridia*, and *Myxococcia* species highlighting a potentially broad role in anti-phage defense (Fig. 1a and Extended Data Fig. 1a). Sequence alignments of the dNTPase proteins demonstrate strict conservation of a canonical HD motif and HE catalytic dyad required for dNTP hydrolysis as observed in histidine-aspartate (HD) domain superfamily proteins(Aravind & Koonin, 1998; Ji et al., 2014; Vorontsov et al., 2011) (Fig. 1b). Likewise, alignments of the accessory NTase protein demonstrate conservation of an active site [D/E]h[D/E]..h[D/E]h motif characteristic of other Pol-β-superfamily immune-related nucleotidyltransferase proteins in CBASS and Hailong anti-phage defense(Cohen et al., 2019; Tan et al., 2025; Whiteley et al., 2019) (Fig. 1b). We cloned representative dNTPase-NTase systems from *Salmonella enterica* SA20044414 and *Escherichia coli* H5, as well as a control dGTPase protein from *Shewanella putrefaciens* CN-32 (*Sp*dGTPase) previously demonstrated to inhibit phage(Tal et al., 2022), and tested the ability of these systems to defend against phage infection. *Salmonella* and *E. coli* dNTPase-NTase systems exhibited strong anti-phage defense with >10,000-fold protection against phage T5 and related T5-like phages in the family *Demerecviridae* but only weakly defended against phage T7 and other distantly related phages(Maffei et al., 2021) (Fig. 1c and Extended Data Fig. 1b). In contrast, *Sp*dGTPase strongly defended against phage T7 but only weakly against phage T5 suggesting a differential ability of dNTPase-NTase systems to protect against phage infection (Fig. 1c and Extended Data Fig. 1b). Mutations to the conserved dNTPase HD motif in the dNTPase-NTPase systems abolished all phage defense verifying that dNTPase function is required for phage inhibition (Fig. 1c and Extended Data Fig. 1b,c). Interestingly, mutations to the active site of the accessory NTase protein induced cellular toxicity with the mutated *E. coli* dNTPase-NTase^DDAA^ operon in particular resulting in >10,000-fold loss of bacterial fitness (Fig. 1d). Together, these results demonstrate dNTPase-NTase operons are anti-phage defense systems and suggest that the accessory NTase protein has a role in suppressing dNTPase-associated toxicity.

**Figure 1.**
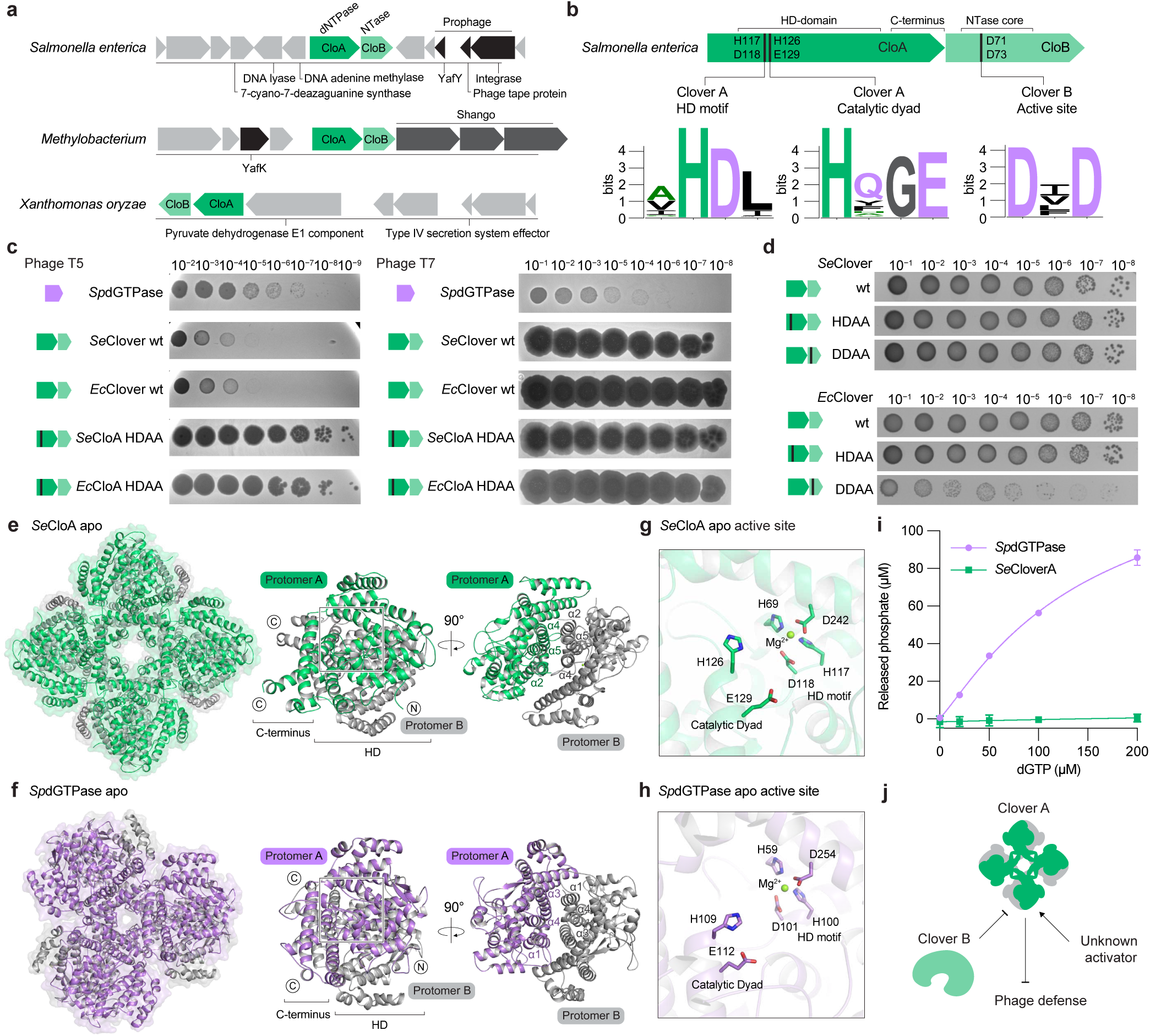
Discovery of Clover anti-phage defense system. **a**, Representative Clover operons and gene neighborhoods from *Salmonella enterica* SA20044414, *Methylobacterium symbioticum* SB0023/3, and *Xanthomonas oryzae* DXO-052. Defense systems were annotated using DefenseFinder(Tesson et al., 2024). **b**, Schematics of CloA and CloB proteins with sequence logos depicting conservation of active site residues. **c**, Representative plaque assays of *Shewanella putrefaciens* CN-32 dGTPase (*Sp*dGTPase), and Clover, CloA catalytic mutant, or CloB catalytic mutant operons from *Salmonella enterica* SA20044414 or *Escherichia coli* H5. **d**, Bacterial growth assay of a 10-fold dilution series of *E. coli* containing arabinose-inducible plasmids expressing *Salmonella enterica* or *Escherichia coli* Clover wild-type operons, CloA HD motif mutant, or CloB inactive mutant (D71A/D73A). **e**, Cryo-EM structure of *Salmonella enterica* SA20044414 CloA (*Se*CloA) and views of the *Se*CloA dimeric unit. **f**, Crystal structure of anti-phage *Sp*dGTPase and views of the *Sp*dGTPase dimeric unit. **g**, Overview of CloA active site including the metal binding HD motif and conserved H126/E129 residues. **h**, Overview of *Sp*dGTPase metal binding HD motif and H126/E129 active site. **i**, Phosphate release assay measuring dGTP hydrolysis by *Se*CloA and *Sp*dGTPase. Purified enzymes were incubated with indicated dGTP concentration and phosphate release was quantified using malachite green. **j**, Schematic of a hypothesized mechanism of action of the Clover defense system.

To define the mechanism of dNTPase-NTase regulation, we determined a 2.4 Å cryo-EM structure of the *Salmonella enterica* dNTPase protein (Fig. 1e, Extended Data Fig. 1d, and Extended Data Fig. 2). The *Salmonella* dNTPase comprises an N-terminal HD domain and a C-terminal α-helical bundle and forms an octameric complex consisting of four pairs of dimers that arrange to create an architecture resembling a four-leaf clover (Fig. 1e). We therefore named the dNTPase-NTase anti-phage defense system Clover and designated the dNTPase protein as Clover protein A (CloA) and the accessory NTase protein as Clover protein B (CloB). The clover-like CloA octameric assembly is formed through extensive interactions across three distinct interfaces. First, HD domain helices α2–α5 interact between adjacent CloA protomers to create anti-parallel dimeric units with α2_A_, α4_A_, and α5_A_ forming primarily hydrophobic interactions with α4_B_, and the α3_A_ residue R46 forming hydrogen-bonding interactions with the peptide backbone of the α3_B_–α4_B_ loop (Fig. 1e and Extended Data Fig. 1e). Second, CloA_AB_ dimer units multimerize through interactions between the CloA_A_ C-terminal α-helical bundle helices α18_C_ and α22_C_ and helices α6_A_ and α17_A_ that interlock the CloA_AC_ dimer (Extended Data Fig. 1f). Third, the CloA_AB_ dimer unit interacts with CloA_D_, forming pi-stacking interactions between α12 residue H263 of CloA_A_ and CloA_D_ thereby stabilizing the CloA_ABCD_ tetramer. (Extended Data Fig. 1g). To further understand the architectural organization of anti-phage defense dNTPase proteins we next determined a 2.7 Å X-ray crystal structure of *Sp*dGTPase(Tal et al., 2022). *Salmonella* CloA and *Sp*dGTPase have low ∼33% overall amino-acid homology but adopt the same overall HD fold and share all major structural features (Fig. 1f and Extended Data Fig. 1h,i). However, unlike the octameric assembly of CloA, alterations in the *Sp*dGTPase multimerization interface result in assembly of a hexameric complex (Fig. 1f and Extended Data Fig. 1j). Similar to CloA, the core unit of *Sp*dGTPase is an anti-parallel dimer formed through hydrophobic interactions throughout the HD domain. Differences in CloA and *Sp*dGTPase overall complex architecture result from changes in the CloA C-terminal α-helical bundle where lengthening of helices α19 and α22 causes the CloA dimeric units to interlock at an angle that is ∼30° wider and multimerize as an octameric assembly (Fig. 1f and Extended Data Fig. 1j).

CloA and *Sp*dGTPase share near identical catalytic and substrate nucleotide binding site residues suggesting that CloA likely also functions as a dGTPase (Extended Data Fig. 1j). Similar to canonical dGTPases like the basal *E. coli* housekeeping enzyme Dgt(Barnes et al., 2019), the CloA HD active site motif H117 and D118 coordinate a catalytic Mg^2+^ ion along with residues H69 and D242, and the HE catalytic dyad H126 and E129 are positioned for substrate dGTP hydrolysis (Fig. 1g,h and Extended Data Fig. 1k). We tested enzyme function *in vitro* and observed that *Sp*dGTPase is alone sufficient to catalyze rapid dGTP hydrolysis (Fig. 1i). Interestingly, in contrast to *Sp*dGTPase, no activity was observed with CloA enzyme alone, suggesting that Clover dGTPase activity requires recognition of a cue that occurs during phage infection. Together our results explain structural changes in anti-phage defense dNTPase proteins that control multimerization and complex assembly and suggest a model of regulation in Clover anti-phage defense where two distinct signals control activation and suppression of CloA dGTPase function (Fig. 1j).

### Clover defense senses increased cellular dTTP

To understand how phage infection triggers activation of Clover anti-phage defense, we infected *E. coli* cells expressing the *Salmonella* Clover system and isolated escape phage mutants. Phage T5, Bas28, and Bas29 escape mutants replicated to high titers with >10,000-fold recovery and exhibited clear resistance to Clover defense (Fig. 2a). Whole genome sequencing of six resistant phages revealed missense and premature nonsense mutations in viral genes encoding deoxyribonucleoside monophosphatase (Dmp) and ribonucleoside diphosphate reductase (subunits NrdA and NrdB) enzymes (Fig. 2b). Both Dmp, a phosphatase that hydrolyzes deoxynucleotides to prevent recycling of host dNTPs, and NrdAB, a reductase that converts ribonucleotides to deoxyribonucleotides, are conserved in all T5-like phages and function to remodel the cellular dNTP pool and support phage DNA replication(Eriksson & Berglund, 1974; Mozer & Warner, 1977; Warner et al., 1975). Mapping escape mutations onto AlphaFold3 predicted structures of Dmp and a previously reported model of *E. coli* NrdAB suggested these mutations likely modulate, but do not ablate, the ability of phages to modify intracellular dNTP levels(Kang et al., 2020) (Extended Data Fig. 3a,b).

**Figure 2.**
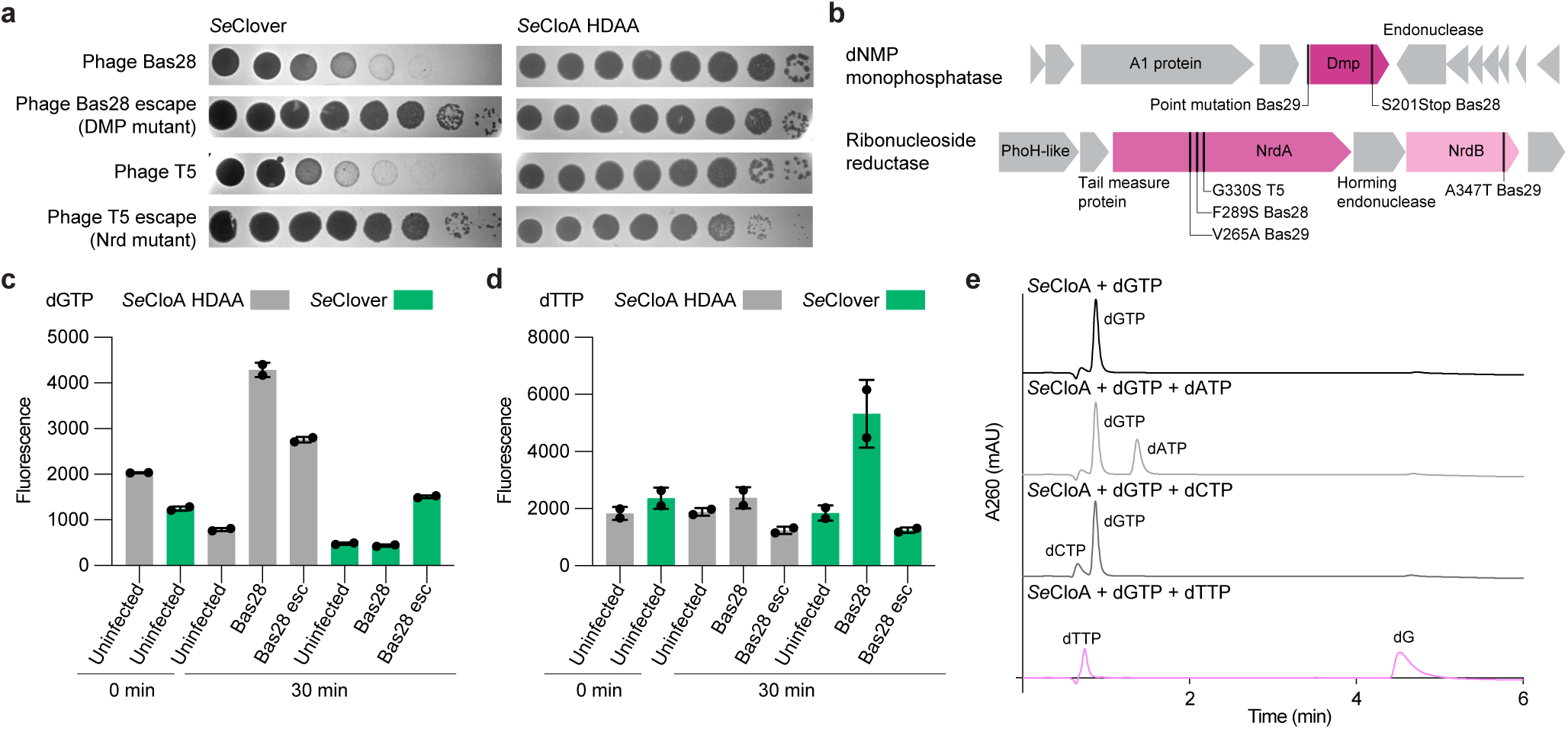
Clover defense senses increased cellular dTTP. **a**, Representative plaque assays of Clover or *Se*CloA catalytic operon mutant expressing cells challenged with Bas28^WT^, T5^WT^, or phage escape mutants Bas28^Dmp S201Stop^ and T5^NrdA G330S^ in a tenfold serial dilution. **b**, Schematic of the mapped phage Bas28 and phage T5 Clover escape mutants and corresponding genomic loci. Clover escape mutants map to the dNMP monophosphatase (*dmp*) and ribonucleotide reductase (*nrdA*, *nrdB*) genes. **c** and **d**, Changes in intracellular dGTP and dTTP concentrations measured in *Salmonella enterica* Clover wild-type and CloA catalytic inactive mutant (H117A/D118A) operons expressing *E. coli* infected with Bas28^WT^ and Bas28^Dmp S201Stop^. **e**, LC-MS analysis of purified *Se*CloA incubated with indicated dNTPs demonstrates that *Se*CloA is a dGTPase activated by the presence of dTTP.

We measured intracellular dNTP levels in *E. coli* expressing Clover defense systems with and without phage infection. In cells expressing a dGTPase inactive mutant CloA^HDAA^, dGTP levels were elevated following infection with wildtype phage Bas28 and phage T5 in agreement with the known ability of T5-like phages to increase intracellular dNTP concentrations(Ahluwalia et al., 2012; Eriksson & Berglund, 1974; Mozer et al., 1977; Tal et al., 2022) (Fig. 2c and Extended Data Fig. 3c). Compared to wildtype phages, the phage Bas28 escape mutant (Bas28^Dmp S201Stop^) exhibited decreased overall accumulation of dGTP and the phage T5 escape mutant (T5^NrdA G330^) exhibited similar levels of dGTP accumulation (Fig. 2c and Extended Data Fig. 3e). In *E. coli* expressing the wildtype Clover system, infection with wildtype phages resulted in a significant reduction in intracellular dGTP levels, suggesting that phage infection triggered activation of Clover defense and induction of the CloA dGTPase (Fig. 2c and Extended Data Fig. 3e). In contrast, in cells expressing Clover and infected with Bas28^DmpS201Stop^, we observed dGTP concentrations similar to the CloA^HDAA^ control, demonstrating that infection with the escape mutant phage resulted in reduced activation of Clover defense (Fig. 2c and Extended Data Fig. 3e). Clover expressing cells infected with wildtype phages additionally exhibited increased accumulation of dTTP consistent with phage enzymes driving elevated dNTP production to compensate for dGTP replete conditions(Tal et al., 2022) (Fig. 2d and Extended Data Fig. 3f). Notably, in cells infected with the phage Bas28^Dmp S201Stop^ escape mutant we no longer observed elevated dTTP concentrations suggesting a connection between elevated intracellular dNTP levels and activation of Clover defense (Fig. 2d and Extended Data Fig. 3f).

To test a potentially direct role for CloA in sensing intracellular dNTP concentrations, we next measured the ability of CloA to respond to dNTPs *in vitro*. Remarkably, CloA reactions supplemented with dTTP exhibit dramatic activation of dGTPase activity and rapidly hydrolyze dGTP substrate (Fig. 2e and Extended Data Fig. 3g). CloA activation is highly specific, with both *Salmonella* and *E. coli* CloA proteins unaffected by dATP or dCTP and responsive only to the presence of elevated concentrations of dTTP (Fig. 2e and Extended Data Fig. 3g). Likewise, CloA exhibits strict dGTP substrate specificity, with the activated enzyme only hydrolyzing dGTP and leaving other dNTPs including dTTP unaffected (Fig. 2e and Extended Data Fig. 3g). Together these results demonstrate that CloA is a dGTPase effector that senses phage infection by specifically responding to elevated dTTP concentrations associated with viral replication.

### The nucleotide signal p3diT restrains Clover defense

Expression of CloA alone in the absence of CloB results in potent cellular toxicity (Fig. 1d). We hypothesized that the CloB NTase enzyme may therefore synthesize a signaling molecule that directly binds to CloA and prevents aberrant activation of Clover defense. To test this hypothesis, we purified *Salmonella enterica* CloA from cells co-expressing CloB and determined a 2.7 Å cryo-EM structure of the resulting CloA complex (Fig. 3a). We observed strong difference density at the interface between each CloA_A_ and CloA_B_ protomer in a pocket lined by highly conserved charged residues, including R29_A_, R34_A_, R38_A_, and R314_B_ (Fig. 3b and Extended Data Fig. 4a–c). We observed strong density that could be unambiguously assigned as a nucleotide ligand composed of thymidine di- or tri-phosphate connected to a second thymidine base with a canonical 3′–5′ phosphodiester linkage, and we named this nucleotide signal p3diT (5′-triphosphothymidyl-3′5′-thymidine) (Fig. 3c and Extended Data Fig. 4a).

**Figure 3.**
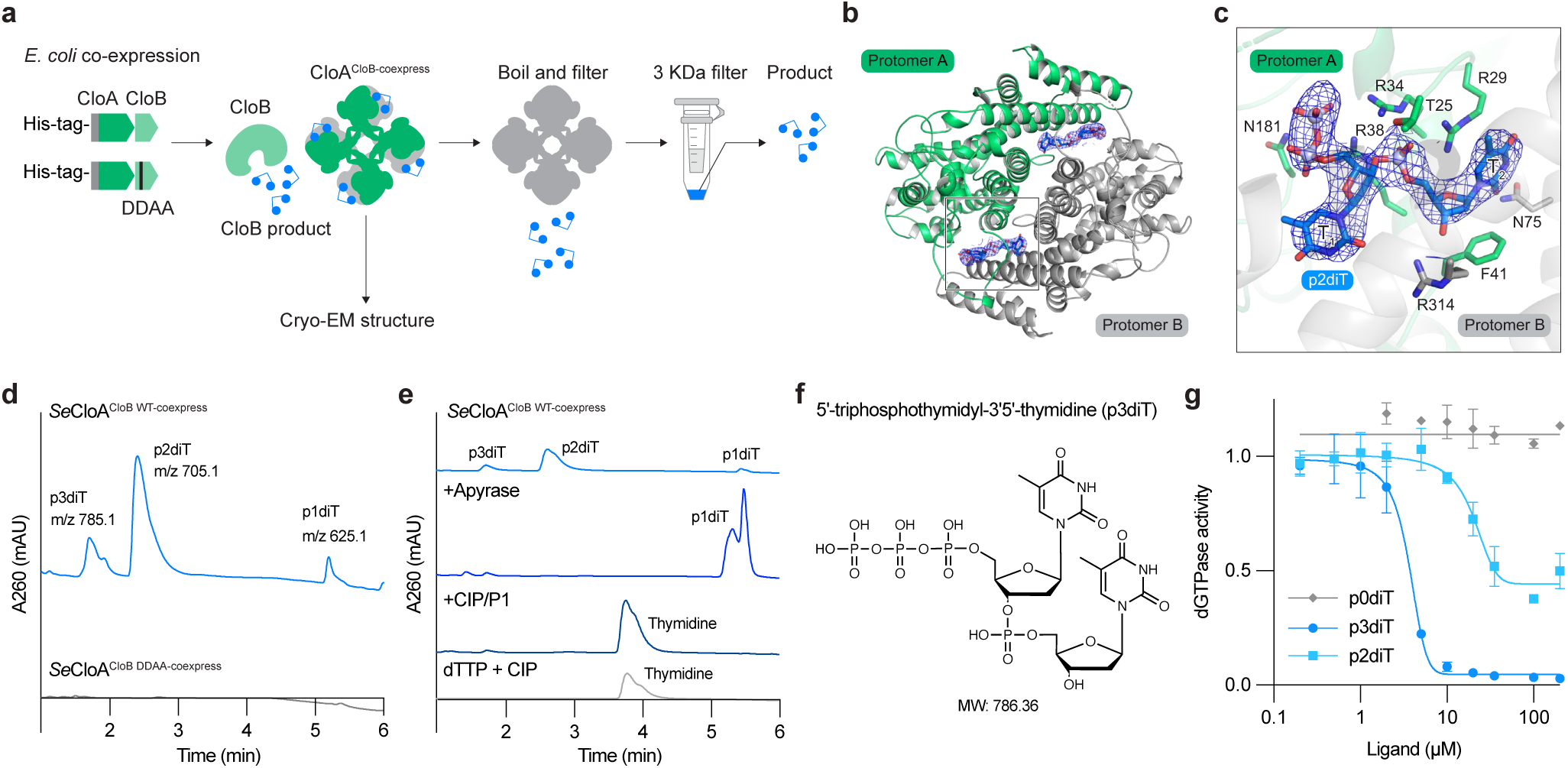
The nucleotide signal p3diT restrains Clover defense. **a**, Schematic representation of strategy used to identify the CloB nucleotide signal from cells co-expressing CloA and CloB. His-tagged *Se*CloA from cells with co-expressed wild-type CloB or CloB catalytic inactive mutant (D61A/D63A) (CloB^DDAA^) was purified. The *Se*CloA complex was then analyzed by cryo-EM structural analysis and in parallel heat denatured to release the CloB signal for direct chemical analysis. **b**, Dimer view of cryo-EM structure of *Se*CloA from cells co-expressing CloB along with the 2F_O_–F_C_ density (blue) of the CloB nucleotide signal. **c**, Detailed view of CloB-derived nucleotide signal interacting within the CloA dimer interface. **d**, LC-MS analysis of boiled *Se*CloA co-expressed with CloB or CloB^DDAA^. **e**, LC-MS analysis of apyrase and CIP/P1 treatment of boiled CloA product, and CIP treatment of dTTP. **f**, Chemical structure of the CloB nucleotide signal 5′-triphosphothymidyl-3′5′-thymidine (p3diT). **g**, Inhibition of *Se*CloA activity by p0diT, p2diT, and p3diT. Wild-type *Se*CloA was incubated with dGTP, dTTP, and either p0diT, p2diT, or p3diT and then phosphate release was quantified using malachite green.

To verify that CloB synthesizes the nucleotide immune signal p3diT, we next isolated p3diT from the purified CloA complex for direct chemical analysis (Fig. 3a,d). Using liquid chromatography-tandem mass spectrometry (LC-MS), we observed ions of peaks of 785.1, 705.1, and 625.1 *m/z*, corresponding to the masses of p3diT and the related hydrolyzed diphosphate (p2diT) and monophosphate (p1diT) forms (Fig. 3d and Extended Data Fig. 4d). No nucleotide product masses were identified when CloA was expressed with a catalytically inactive CloB mutant D71A/D73A enzyme, supporting that CloB is responsible for synthesis of p3diT (Fig. 3d and Extended Data Fig. 4d). To verify the 5′ phosphorylation status of p3diT, we treated the purified nucleotide signal with apyrase, a phosphatase that selectively removes 5′ γ and β phosphates, and confirmed complete conversion of p3diT to p1diT (Fig. 3e and Extended Data Fig. 4e). Likewise, to verify the thymidine base identity of p3diT we digested the nucleotide signal with nuclease P1 and alkaline phosphatase and confirmed complete conversion of p3diT to a product with mass corresponding to thymidine (Fig. 3e,f and Extended Data Fig. 4f). Finally, we verified that the mass and retention time of the CloA-bound product coincided with a chemically synthesized standard confirming that the nucleotide immune signal in Clover defense is p3diT (Extended Data Fig. 4g).

We next tested the effect of p3diT on CloA dGTPase activity. Introduction of p3diT was alone sufficient to inhibit all CloA dGTPase activity *in vitro* with an IC_50_ of ∼7.6 μM (Fig. 3g). We observed that p2diT was less efficient at inhibiting CloA dGTPase activity (IC_50_ of ∼20.9 μM) and that dephosphorylated p3diT (p0diT) had no detectable effect, consistent with the 5′ terminal phosphates of p3diT being required for high-affinity interaction with the charged CloA ligand binding pocket (Fig. 3g). We repeated CloB product purification experiments with a distantly related CloB enzyme from *Xanthomonas* and confirmed that p3diT signaling is broadly conserved in Clover defense (Extended Data Fig. 4h). These results reveal p3diT as a novel nucleotide immune signal that negatively regulates Clover anti-phage defense.

### Mechanism of nucleotide signal activation and inhibition

To define the mechanism of nucleotide signaling in Clover anti-phage defense, we mutated the *Salmonella* CloA active site (H126A/E129A) to allow direct comparison of dGTP substrate interactions and determined cryo-EM structures of CloA in the suppressed CloA–dGTP–p3diT (2.6 Å) or activated CloA–dGTP–dTTP (2.6 Å) states (Fig. 4a, Extended Data Fig. 6a). Remarkably, the CloA–dGTP–p3diT and CloA–dGTP–dTTP structures reveal that p3diT and dTTP each bind at the CloA_AB_ interface in structurally distinct sites that remodel to accommodate the inhibiting or activating nucleotide signals (Fig. 4a–d). In the CloA–dGTP– p3diT suppressed state structure, p3diT is bound at the CloA_AB_ interface in a pocket formed by helices α2_A_, α4_B_, α14_B_, and loop α1_A_–α2_A_ that seals around the inhibitory nucleotide signal (Fig. 4a, Extended Data Fig. 6a). Conserved residues from each CloA protomer form 10 hydrogen-bond interactions with p3diT that coordinate both the nucleotide signal 5′ triphosphate (CloA R34_A_, R38_A_, and N181_A_) and 3′–5′ phosphodiester linkage (R29_A_, R34_A_, and T25_A_) (Fig. 4a,c). Specificity for p3diT binding is mediated by CloA residues that make base-specific contacts with the Watson-Crick edges of the first thymidine base T_1_ (Q311_B_) and second thymidine base T_2_ (D37_A_ and N75_B_). CloA residues R314_B_ and F41_A_ further read-out the 3′ OH and deoxyribose sugar backbone of base T_2_ (Fig. 4a,c). In the CloA–dGTP–dTTP active state structure, dTTP is bound at the CloA_AB_ interface in an adjacent pocket to the p3diT binding site formed by loops α3_A_–α4_A_ and α3_B_–α4_B_ from each CloA protomer and the C-terminal helical bundle α18_B_–α21_B_ (Fig. 4c, Extended Data Fig. 6b). A distinct set of conserved CloA residues form 13 hydrogen-bond interactions with dTTP that coordinate the nucleotide triphosphate moiety at each of the α-(R47_A_ and S418_A_), β-(K256_B_), and γ-phosphate (R47_A_, Q172_A_, and N420_A_) positions (Fig. 4d). The dTTP ribose sugar backbone is coordinated by the side-chain of CloA residue R47_A_ and the peptide main-chain of residue A444_A_ (Fig. 4d). The thymidine base is recognized by an extensive series of interactions with CloA residues R47_A_ and Y452_A_ that sandwich the nucleobase face and residues D61_B_ and T449_A_ that make sequence-specific contacts along the Watson-Crick edge and explain the exquisite specificity of CloA for dTTP as an activating nucleotide signal (Fig. 4d).

**Figure 4.**
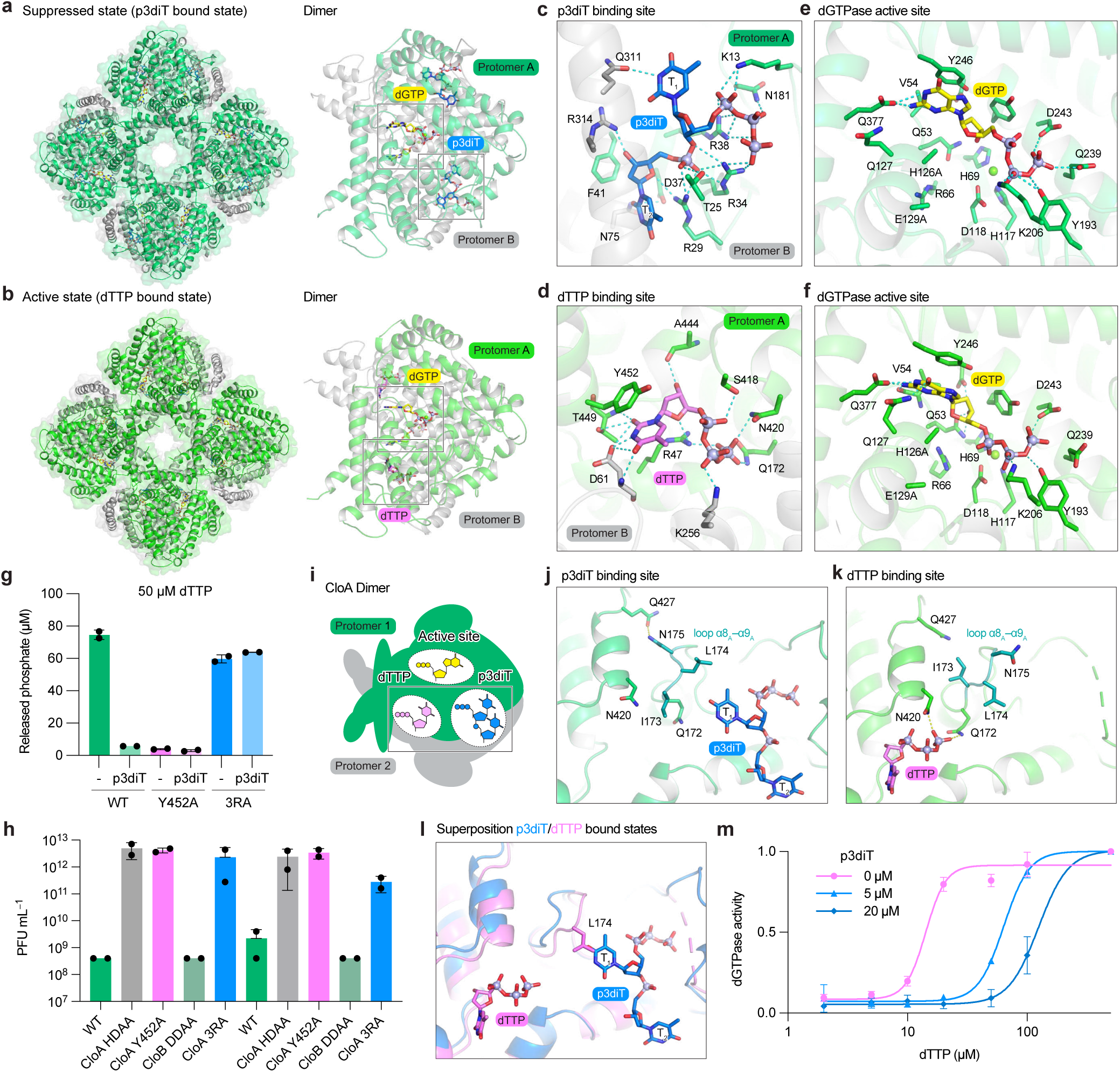
Mechanism of nucleotide signal activation and inhibition. **a**, Cryo-EM structure of *Se*CloA catalytic inactive mutant (H126A/E129A) (CloA^HEAA^) in complex with dGTP and p3diT. dGTP binds within the active site and p3diT binds at the CloA dimeric interface. **b**, Cryo-EM structure of *Se*CloA^HEAA^ in complex with dGTP and dTTP. dGTP binds within the active site and dTTP binding at a distinct pocket from p3diT at the CloA dimeric interface. **C** and **d**, Magnified views of the *Se*CloA p3diT and dTTP binding pockets. **e** and **f**, Magnified views of dGTP active site contacts in the *Se*CloA–dGTP–p3diT and *Se*CloA–dGTP– dTTP complex structures demonstrates that a dTTP-induced conformational change completes the CloA active site and repositions the dGTP substrate for hydrolysis. **g**, Phosphate release assay measuring dGTP hydrolysis by wild type *Se*CloA and *Se*CloA dTTP or p3diT binding mutants (Y452A and 3RA) with or without p3diT. Bar graph depicts the mean, with error bars representing standard deviation. **h**, Effect of p3diT and dTTP binding pocket mutations on Clover anti-phage defense. Data represent plaque-forming units per mL (PFU mL^−1^) of T5 phages. Bar graph depicts the mean, with error bars representing standard deviation. **i**, Cartoon schematic of the CloA dimeric unit depicting distinct active site, p3diT, dTTP binding pockets. **j** and **k**, Magnified views of p3diT and dTTP binding pocket as in (**i**) in p3diT bound and dTTP bound states demonstrates a key conformational change in CloA loop α8_A_–α9_A_. **l**, Superposition of *Se*CloA–dGTP–p3diT and *Se*CloA–dGTP–dTTP complex structures demonstrates the conformation change that occurs between p3diT and dTTP binding and a key role for CloA L174_A_ on loop α8_A_–α9_A_ that sterically occludes p3diT in the dTTP-bound state. **m**, Activation of *Se*CloA activity by dTTP in the presence of p3diT. Wild-type *Se*CloA incubated with dGTP and different concentrations of dTTP with or without p3diT.

Superposition of the dGTPase active site from the CloA–dGTP–p3diT and CloA–dGTP– dTTP structures reveals why the nucleotide signal dTTP is required to enable dGTP hydrolysis (Fig. 4e,f). dGTP is present in both the CloA active and suppressed state structures, demonstrating that nucleotide signaling is not required for initial substrate recognition (Fig. 4a,b). However, in the p3diT-bound suppressed state, the dGTP substrate is coordinated by comparatively minimal interactions between CloA residues that contact the guanosine nucleobase (Y246_A_ and Q377_A_) and β- and γ-phosphates (Y193_A_, D243_A_, and K206_A_) (Fig. 4e). In contrast, in the dTTP-bound active state, dGTP is coordinated by an additional set of five contacts that reorient substrate interactions and complete active site formation (Fig. 4f). First, dTTP-binding induces a conformational change that repositions sidechain R66_A_ to form new hydrogen bond interactions with the peptide backbone of T52_A_, Q53_A_, and N121_A_ that stabilize loop α3_A_–α4_A_ in the dGTPase core (Extended Data Fig. 6c–e). Stabilization of CloA loop α3_A_– α4_A_ completes the dGTPase active site and creates additional interactions with dGTP including contacts to the guanosine base (sidechain of Q127_A_ and mainchain of Q53_A_ and V54_A_) and the 3′ OH (sidechain of Q53_A_) (Fig. 4f and Extended Data Fig. 6e). Second, substrate reorientation in the CloA–dGTP–dTTP active state structure positions the dGTP α-phosphate moiety in proximity to the H126A/E129A catalytic dyad in a conformation that is now competent for dGTP hydrolysis (Extended Data Fig. 6f). We introduced a CloA Y452A mutation to disrupt the dTTP-binding site and observed loss of all dGTP hydrolysis activity in the presence of elevated dTTP *in vitro* and elimination of Clover anti-phage defense *in vivo* (Fig. 4g,h and Extended Data Fig. 6g).

To explain how p3diT signaling regulates CloA dGTPase activity, we next compared the CloA_AB_ interfaces in the CloA–dGTP–p3diT and CloA–dGTP–dTTP structures (Fig. 4i–l). In the p3diT bound suppressed state, interactions with the inhibitory nucleotide signal stabilize CloA loop α8_A_–α9_A_ in an open conformation that is drawn away from the dTTP-binding site (Fig. 4j). Within CloA loop α8_A_–α9_A_, sidechain N175_A_ hydrogen bonds with sidechain Q427_A_ to stabilize this open conformation and reorient CloA residues Q172_A_ and N420_A_ away from the dTTP-binding site to disfavor interactions with the activating signal (Fig. 4j,k). Superposition of the p3diT- and dTTP-bound structures demonstrates a clash between p3diT in the CloA suppressed structure and residue L174_A_ in the CloA active state structure, further suggesting that p3diT binding and dTTP binding are mutually exclusive (Fig. 4l). To test this hypothesis, we first introduced CloA mutations to conserved arginine residues in the p3diT binding-site (R29A/R34A/R38A, 3RA) to selectively disrupt interactions with the inhibitory nucleotide signal. The CloA 3RA mutant exhibited robust dGTPase activity *in vitro* but was no longer inhibited by the presence of p3diT and was sufficient to coordinate productive regulation of Clover defense *in vivo* (Fig. 4g,h and Extended Data Fig. 6g,h). Next, we examined the concentration of dTTP required to activate CloA dGTPase activity in the presence or absence of p3diT regulation. In the absence of p3diT regulation, dTTP potently induces CloA dGTPase activity and exhibits an EC_50_ of ∼15 μM (Fig. 4m). Consistent with our structural comparisons, introduction of increasing concentrations of p3diT limits CloA activation and necessitates a progressively higher level of the activating signal dTTP to induce dGTPase function, with >200 μM of dTTP required to overcome 20 μM of p3diT signal (Fig. 4l,m). Together, these results reveal the molecular basis of Clover nucleotide signal regulation and explain how activating and inhibiting nucleotide signals can cooperatively control bacterial anti-phage defense.

Our study defines Clover as a bacterial immune system that surveys and targets the cellular nucleotide pool to defend against phage infection. Clover defense uses a remarkable series of signals that connect cellular nucleotide availability to each aspect of antiviral immunity, including viral sensing, restriction of viral replication, and protection from immune-associated toxicity. To detect infection, the Clover protein CloA senses elevated dTTP concentrations as a molecular cue associated with phage replication. In addition to known mechanisms that sense viral protein or nucleic acid(Banh et al., 2023; Burman et al., 2024; Deep et al., 2024; Gao et al., 2022; Garb et al., 2022; Haudiquet et al., 2025; Jaskólska et al., 2022; Kibby et al., 2024; Loeff et al., 2025; Pradhan et al., 2024; Robins et al., 2024; Roisné-Hamelin et al., 2024; Zhang et al., 2022), Clover defense reveals that bacterial immunity can monitor cellular metabolite pools to guard against phage infection. Following nucleotide detection and CloA–dTTP complex formation, a conformational change activates CloA dGTPase effector function to deplete the cellular nucleotide pool and restrict phage replication. Clover defense encodes an accessory enzyme CloB that synthesizes an additional nucleotide signal p3diT that suppresses CloA activation and protects uninfected cells from aberrant nucleotide depletion. p3diT binds to CloA in an allosteric site adjacent to the dTTP binding pocket and raises the threshold of intracellular dNTP levels required to trigger full Clover immune activation. Bacterial, plant, and animal cells are known to synthesize an enormous diversity of nucleotide immune signals to control immune activation(Athukoralage & White, 2022; Hobbs & Kranzusch, 2024; Maruta et al., 2023; Slavik & Kranzusch, 2023). Clover p3diT joins two recently described nucleotide signals oligodeoxyadenylate in Hailong defense and 2′3′-c-di-AMP in MISS/Panoptes defense that surprisingly invert this paradigm and negatively regulate immune activation(Doherty et al., 2025; Sullivan et al., 2025; Tan et al., 2025). Together, our findings define a mechanism of Clover anti-phage defense and discover how distinct nucleotide signals cooperate to coordinate activation and inhibition of antiviral immunity.

## Supporting information

Extanded Figure

## Methods

### Bioinformatics analysis of CloB

The genomic sequences of CloB and closely related family members were aligned using MAFFT as previously described(Tan et al., 2025). Briefly, sequences were identified using IMG BLAST searches performed with an E-value cut-off of 0.005, with CloB from *Salmonella* (IMG ID 2897288308) and *Clostridioides difficile* VRECD0136 (IMG ID 2904220186) as query proteins, and duplicates were removed. Clover operon genetic neighborhoods were manually curated from IMG and annotated using Defense Finder(Tesson et al., 2024). Sequences of CloB homologs are reported in Supplementary Table 1.

### Bacterial strains and growth conditions

*E. coli* strain BW25113 was grown in LB medium (1% tryptone, 0.5% yeast extract, and 1% NaCl w/v) supplemented with 0.1 mM MnCl_2_, 5 mM MgCl_2_, 5 mM CaCl_2_ with or without 0.5% agar) at 37°C or room temperature (RT). Whenever applicable, media were supplemented with ampicillin (100 µg mL^−1^) and/or chloramphenicol (34 µg mL^−1^) to ensure the maintenance of plasmids. *E. coli* strain BL21-RIL (Agilent) was used for all protein expression experiments and *E. coli* strain DH10β (strain Top10, Invitrogen) was used for cloning and plasmid propagation. For repression of protein expression from pET vectors, BL21 *E. coli* was cultivated in MDG medium (0.5% glucose, 25 mM Na_2_HPO_4_, 25 mM KH_2_PO_4_, 50 mM NH_4_Cl, 5 mM Na_2_SO_4_, 2 mM MgSO_4_, 0.25% aspartic acid and trace metals) with ampicillin and chloramphenicol. For optimal protein expression from pET vectors, BL21 *E. coli* was cultivated in M9ZB medium (0.5% glycerol, 1% Cas-amino acids, 47.8 mM Na_2_HPO_4_, 22 mM KH_2_PO_4_, 18.7 mM NH_4_Cl, 85.6 mM NaCl, 2 mM MgSO_4,_ and trace metals) supplemented with ampicillin and chloramphenicol.

### Cloning and plasmid construction

For protein purification, genes of *CloA* from *Salmonella* (IMG ID 2897288307) and *E. coli* H5 (IMG ID 2719124844), and *Shewanella putrefaciens dGTPase* were synthesized as gBlocks (Integrated DNA Technologies), and cloned into custom pET vectors with an N-terminal 6×His-SUMO2-5×GS tag fusion(Zhou et al., 2018). Genes of *CloB* from *Salmonella* (IMG ID 2897288306), *E. coli* (IMG ID 2719124845), and *Xanthomonas oryzae* DXO-052 (IMG ID 8060537493) were synthesized as gBlocks (Integrated DNA Technologies) and cloned downstream of *CloA* sequence with a ribosomal binding site.

For phage challenge, bacterial toxicity, and dNTP quantification assays, the Clover operon from *Salmonella* (IMG ID 2897288195 104388–106590) and *E. coli* H5 (IMG ID 2719124844 82910–85109), and *Shewanella putrefaciens* dGTPase was synthesized (Twist Bioscience) and cloned into a pBAD arabinose-inducible vector(Hobbs et al., 2022). Mutations and deletions were introduced using site-directed mutagenesis PCR, and the sequences were confirmed by whole-plasmid sequencing (Plasmidsaurus).

## Phage challenge assay

Liquid cultures were inoculated from single bacterial colonies and incubated with shaking at 37°C for 4 h until OD_600_ reached at least 0.6. Cultures were diluted to an OD_600_ of 0.3 in melted top agar supplemented with L-arabinose and then poured onto a bacterial plate containing 0.02% (w/v) L-arabinose. The plate was then allowed to solidify at room temperature for 2 h. Phages were prepared in 10-fold serial dilutions from high-tier stocks using SM buffer (50 mM Tris-HCl pH 7.5, 100 mM NaCl, 8 mM MgSO_4_) and spotted as 2.5 μL drops onto the cooled top agar. Plates were incubated at room temperature for 30 min to dry and incubated overnight at 30°C. Plates were imaged using a Bio-Rad ChemiDoc MP, and phage plaques were manually counted from the images. In cases where individual phage plaques could not be distinguished, the highest dilution phage drop with observable bacterial clearance was counted as 10 plaques.

### Bacterial toxicity assay

Liquid cultures were inoculated from single bacterial colonies and incubated with shaking at 37°C overnight. Bacteria were pelleted by centrifugation and resuspended in phosphate-buffered saline (PBS). 10-fold serial dilutions were performed with PBS, and diluted bacteria were spotted in 5 μL drops onto plates supplemented with 0.02% L-arabinose or 1% D-glucose. Plates were dried at room temperature for 30 min and incubated overnight at 30°C. Plates were imaged using a Bio-Rad ChemiDoc MP.

### Protein expression and purification

Recombinant wild type *Salmonella* or *E. coli* H5 CloA, *Salmonella* CloA mutants (H126A/E129A, R29A/R34A/R38A, Y452A), and *Salmonella* or *E. coli* H5 co-expressed with *Salmonella, E. coli* H5, and *Xanthomonas* CloB were purified as previously described(Zhou et al., 2018). Briefly, all proteins were cloned into custom pET vector as an N-terminal His-SUMO fusion-tagged protein with a 5×GS linker and transformed into *E. coli* strain BL21-RIL. Large scale cultures (2–4 liters) were grown for 5 h at 37°C, then induced with IPTG overnight at 16°C. Bacterial pellets were resuspended and sonicated in lysis buffer (20 mM HEPES-KOH pH 7.5, 400 mM NaCl, 30 mM imidazole, and 10% glycerol) and purified using Ni-NTA resin (Qiagen). The Ni-NTA resin was washed with lysis buffer supplemented with 1 M NaCl and then eluted with lysis buffer supplemented with 300 mM imidazole. The Ni-NTA elution fraction was dialyzed into 20 mM HEPES-KOH pH 7.5, 250 mM KCl overnight while removing the SUMO2 tag with recombinant human SENP2 protease (D364–L589, M497A). All CloA proteins were concentrated using a 30 K-cutoff concentrator (Millipore) and purified by size exclusion chromatography on a S200 16/60 Sephacryl 300 column (Cytiva). Proteins were concentrated to greater than 10 mg mL^−1^, flash-frozen with liquid nitrogen, and stored at −80°C.

### Cryo-electron microscopy data collection

A concentration of 1 mg mL^−1^ *Salmonella* CloA or CloA expressed with *Salmonella* CloB was applied to glow-discharged grids and vitrified in liquid ethane on TEM grids using a Mark IV Vitrobot (ThermoFisher). A 3 µL solution of each sample was independently deposited onto a 2/1 Au Quantifoil grid with Carbon mesh, then blotted with filter paper for 4 seconds, using a double-sided blot with a force of 10, in a 100% relative humidity chamber at 4°C. *Salmonella* CloA apo, CloA co-expressed with CloB, CloA^HEAA^–dGTP–p3diT, and CloA^HEAA^–dGTP–dTTP complex were screened and imaged using a Talos Arctica (ThermoFisher) microscope operating at 200 kV and equipped with K3 direct electron detector (Gatan). We collected 6,001 movies of apo *Salmonella* CloA and 3,146 movies of *Salmonella* CloA co-expressed with *Salmonella* CloB on a Titan Krios G3i microscope operated at 300LkV and equipped with a Falcon 4i direct electron detector (Thermo Fisher) with an energy filter. CloA apo and *Salmonella* CloA co-expressed with *Salmonella* CloB data were acquired using EPU software at a pixel size of 0.73 Å and 0.74 Å, respectively. A defocus range of −0.8 to −2.0 µm and a total dose of 49.66 e^−1^ Å was used for *Salmonella CloA* apo, and a total dose of 48.69 e^−1^ Å was used for *Salmonella* CloA co-expressed with *Salmonella* CloB. We collected 3,121 movies of *Salmonella* CloA^HEAA^–dGTP–p3diT or 1,921 movies of *Salmonella* CloA^HEAA^–dGTP–dTTP on a Talos Arctica (ThermoFisher) microscope operating at 200 kV and equipped with K3 direct electron detector (Gatan). *Salmonella* CloA apo and CloA co-expressed with *Salmonella* CloB were acquired using EPU software at a pixel size of 1.1 Å with a defocus range of −0.9 to −2.2 µm and a total dose of 50.24 e^−1^ Å was used for CloA^HEAA^–dGTP–p3diT, and a total dose of 50.46 e^−1^ Å was used for CloA^HEAA^–dGTP–dTTP.

### Cryo-EM data processing

All data processing software was compiled and configured by SBGrid(Morin et al., 2013). Movies were imported into cryoSPARC v.4.4.1 to v.4.6.0(Punjani et al., 2017). Imported movies were processed with patch-based motion correction, patch-based CTF estimation, two-dimensional and three-dimensional particle classification, and homogenous refinement in C_4_ symmetry. The cryoSPARC data-processing procedure is outlined in Extended Data Figs. 3, 5, and 6. In brief, after patch-based CTF estimation, 200 or 1,000 micrographs were selected and autopicked using Blob Picker, and two-dimensional classifications were then used to generate templates for Template Picker. 886,806 particles for the *Salmonella* CloA apo structure, 177,600 particles for the *Salmonella* CloA co-expressed with *Salmonella* CloB, 1,498,407 particles for the *Salmonella* CloA^HEAA^–dGTP–p3diT, and 473,256 particles for the *Salmonella* CloA^HEAA^– dGTP–dTTP were picked. After one round of 2D classification, 324,832 particles for the *Salmonella* CloA apo structure, 221,450 particles for the *Salmonella* CloA co-expressed with CloB, 846,498 particles for the *Salmonella* CloA^HEAA^–dGTP–p3diT, and 312,310 particles for the *Salmonella* CloA^HEAA^–dGTP–dTTP were used in ab initio (KL=L3), followed by heterogeneous refinement, resulting in 2.37 Å for the *Salmonella* CloA apo structure, 2.65 Å for the *Salmonella* CloA co-expressed with CloB, 2.56 Å for the *Salmonella* CloA^HEAA^–dGTP–p3diT, and 2.59 Å for the *Salmonella* CloA^HEAA^–dGTP–dTTP.

### Cryo-EM model building

The starting model, an AlphaFold2-predicted structure of the *Salmonella* CloA octamer, was docked into the electron microscopy density in Coot. P3diT, p2diT, dGTP, and dTTP were generated by Phenix eLBOW option and fitted manually. The model was iteratively refined in Phenix (with non-crystallographic symmetry constraints applied to protein and DNA chains) and manually adjusted in Coot. Structure stereochemistry statistics are reported in Supplementary Table 2. Figures were prepared in PyMOL 2.5.5 (Schrödinger, LLC).

### Crystallization and structure determination

Crystals of *Shewanella putrefaciens* dGTPase were grown in a hanging-drop format using the hanging-drop vapor diffusion method for 3–5 days at 18°C. Recombinant *Shewanella putrefaciens* dGTPase was diluted to 10 mg mL^−1^ in a buffer containing 20 mM HEPES-KOH pH 7.5 and 100 mM KCl. The resultant protein mixture was allowed to equilibrate to 18°C for 10 min and crystals were grown in 96 well trays containing 70 µL reservoir solution and 0.2 µL drops. Drops were mixed 1:1 with purified protein and reservoir solution (0.2 M magnesium chloride, 0.1 M HEPES-NaOH pH 7.5, 30% PEG-400). Crystals were cryo-protected with a reservoir solution supplemented with 10% ethylene glycerol and harvested by flash-freezing in liquid nitrogen.

X-ray diffraction data were collected at Advanced Photon Source Argonne National Laboratory (APS). Data were processed with XDS and Aimless(Evans & Murshudov, 2013; Kabsch, 2010). Experimental phase information was determined by molecular replacement using Phaser-MR in Phenix and a model of *Shewanella putrefaciens* dGTPase structure generated using AlphaFold3(Abramson et al., 2024; McCoy et al., 2007). Model building was performed using Coot, and refinement was carried out using Phenix. A summary of crystallographic statistics is provided in Supplementary Table 3. All structural figures were generated using PyMOL (Version 2.5.4, Schrödinger, LLC).

### dGTPase activity test

Purified 0.5 µM dGTPase with substrate dGTP or ligands was incubated for 20 min with 50 mM KCl, 50 mM HEPES-KOH pH 7.5, 5 mM MgCl_2_, 0.1 mM MnCl_2,_ and 1 U pyrophosphatase (PPase) (New England Biolabs) incubated in 37°C. The reaction was quenched by 1 mM EDTA. *Salmonella* CloA and *Shewanella putrefaciens* dGTPase were incubated with dGTP (0–200 µM) (New England Biolabs) in Figure 1i. Wild type *Salmonella* CloA was incubated with dGTP (100 µM) and p3diT (Biolog), p2diT (Biolog), p0diT (0–100 µM) (Dharmacon) in Figure 3g. Wild type *Salmonella* CloA and *Salmonella* CloA mutants (Y452A and 3RA) were incubated with dGTP (100 µM), dTTP (50 or 200 µM), and with or without p3diT (20 µM) in Figure 4g and Extended Data Figure 6g. Wild type *Salmonella* CloA was incubated with dGTP (100 µM), dTTP (0–500 µM), and with or without p3diT (0, 5, or 20 µM). QuantiChrom^TM^ ATPase/GTPase Assay Kit (DATG-200) was used to quantify phosphate released by dNTP hydrolysis, following the manufacturer’s instructions. Samples were diluted 1:7.5 to ensure released phosphate was within the linear range of detection based on phosphate standards. Absorbance at 650 nm of a ‘‘blank’’ sample with no protein was subtracted from all samples, and then the absorbance was used to calculate the molarity of phosphate released based on a standard curve created using phosphate standards. Samples were measured in technical duplicate and are representative of three independent biological replicates.

In Figure 2e, wild type *Salmonella* CloA (0.5 µM) was incubated with dGTP (200 µM) and either 200 µM dATP (New England Biolabs), dCTP (New England Biolabs), and dTTP (New England Biolabs). The reaction was quenched with 1 mM EDTA and then filtered using a 3K-cutoff concentrator (Amicon Ultra-0.5 mL, Merck). Reactions were analyzed on an Agilent 1260 HPLC equipped with a diode array detector and an Agilent 6125 single quadrupole mass spectrometer in negative ion mode using a reverse-phase Agilent InfinitiLab Poroshell SB-Aq column (2.7-µm particle size, 2.1-mm inner diameter, 100-mm length) at a flow rate of 0.45[mL[min^−1^ with a gradient from 100% solvent A (0.1% ammonium formate) to 100% solvent B (methanol). Data are representative of two independent biological replicates.

### Escape phage isolation

To isolate mutant phages that overcome defense, a drop assay was performed as described above using Clover with the bacterium *E. coli* strain BW25113 expressing either the *Salmonella* Clover operon or CloA catalytic operon mutant (H117A/D118A). Tenfold serial dilutions in SM buffer were prepared for the ancestor phage lysate, and then 2.5 µL drops from the dilution series were dropped on top of the solidified layer containing the bacteria. Single plaques were isolated on the defense-containing strain into 50 μL of SM buffer. Overnight cultures of *E. coli* strain BW25113 harbouring the *Salmonella* Clover operon, were diluted 1:100 in LB supplemented with 0.1 mM MnCl_2_, 5 mM MgCl_2_, 5 mM CaCl_2_ with or without 0.5% agar) and grown at 37°C for 2 hr until reached OD 0.3. We then infected isolated phage with 3 mL culture at OD 0.3 and grown at 37°C overnight. Whole genomes of the escape phages were sequenced by Illumina sequencing. Paired-end reads were aligned to reference genomes (T5 AY543070.1 and Bas28 NC_105115) using Bowtie2(Langmead & Salzberg, 2012). We identified mutations using bfctools (Li, 2011) assuming a ploidy of 1 and default settings. Mutations were filtered for those for which there was at least 150x read depth. Mutations that were present in the escape phage samples and not in the parental genomes were identified as true escape mutations.

### Extraction and measurement of cellular dNTPs

Overnight cultures of BW25113 *E. coli* harboring the *Salmonella* Clover operon and CloA catalytic inactive mutant operon (H117A/D118A), were diluted 1:100 in 50 µL MMB medium (0.02% L-arabinose) and grown at 37°C for 2 hr until culture density reached around OD_600_ of 0.5–0.6. The cultures were diluted to OD_600_ 0.3 and infected with wild type Bas28 phage or escape mutants at a final MOI of 2, and wild type T5 phage or escape mutant at a final MOI of 5. Following the addition of phage, at 0 and 30 min post-infection, 1 mL samples were taken and centrifuged for 5 min at 15,000×g. Pellets were flash-frozen using dry ice and ethanol. The cultured cells were resuspended in 60% MeOH and further denatured by incubation at 95°C for 3 min to quench residual enzymatic activity and facilitate extraction. Supernatants from centrifugation were passed through 3K-cutoff centrifugal filters (Amicon Ultra-0.5 mL, Merck) to remove remaining macromolecules. Residual MeOH was evaporated using a SpeedVac Plus SC110A centrifugal vacuum evaporator (Eppendorf), and the dried pellet was resuspended in 40 µL RNase-free water (Avantor). Cellular dGTP or dTTP levels were quantified using the EvaGreen dNTP quantification method(Purhonen et al., 2020). The cellular extracts were subjected to a Q5 DNA polymerase and EvaGreen-based assay for dGTP and dTTP, using dNTP-specific 197-nucleotide templates (dGTP detection template 5′-GGAGTGAGTGTGAGGTGAATGATGAGTGAGTGTGAGGTGAATGTAGAGTGAGTGTGAGGTGAATGATGAGTGAGTGTGAGGTGAATGTAGAGTGAGTGTGAGGTGAATGATGAGTGAGTGTGAGGTGAATGTAGAGTGAGTGTGAGGTGAATGATGAGTGAGTGTGAGGTGAATGGTTTCTTTGGCGGTGGAGGCGG-3′; dTTP detection template 5′-TCGCTCGCTCTTGCCTCGGTCCTCGCTCGCTCTTGCCTCGGTCCTCGCTCGCTCTTGCCTCGGTCCTCGCTCGCTCTTGCCTCGGTCCTCGCTCGCTCTTGCCTCGGTCCTCGCTCGCTCTTGCCT CGGTCCTCGCTCGCTCTTGCCTCGGTCCTCGCTCGCTCTTGCCTCGGTCCTTTATTTGGCGGTGGAGGCGG-3′), as previously described(Purhonen et al., 2020). The reaction concentrations were 1× Q5 reaction buffer, 0.275 μM primer and 0.25 μM template, 50 μM non-limiting dNTPs, and 1.25 μM EvaGreen (Biotum). The Q5 DNA polymerase (New England Biolabs) concentration was 20 U mL^−1^ for dTTP and 10 U mL^−1^ for dGTP detection. The reaction components were prepared as a 2× master mix, and 5 μL of sample for dGTP detection, and 1 μL of sample and 4 μL of RNase-free water for dTTP detection, were pipetted into a 96-well PCR plate (Thermo Fisher). qPCR was programmed as previously described(Purhonen et al., 2020), and baseline and endpoint fluorescence were read at a temperature above the primer annealing temperature. For the highest sensitivity, the polymerization reaction time at 66 °C was limited to 40 min for dTTP and 20 min for dGTP detection. Samples were measured in technical duplicate and are representative of three independent biological replicates.

### Analysis of CloA and CloB products by LC-MS

Protein stocks at 500 µM of *Salmonella* CloA co-expressed with wild type CloB or the CloB catalytically inactive (D71A/D73A) mutant were denatured by incubation at 95°C for 5 minutes. The CloA bound product was then separated by centrifugation through a 3K-cutoff concentrator (Amicon Ultra-0.5 mL, Merck). The flowthrough was treated with nuclease P1 (Sigma) and Quick CIP phosphatase (New England Biolabs) at 37°C for 30 min, and Apyrase (New England Biolabs) at 30°C for 60 min prior to LC-MS analysis. The dTTP standard was is treated with Quick CIP phosphatase at 37°C for 30 min. Samples were analyzed on an Agilent 1260 HPLC equipped with a diode array detector and an Agilent 6125 single quadrupole mass spectrometer in negative ion mode using a reverse-phase Agilent InfinitiLab Poroshell SB-Aq column (2.7-µm particle size, 2.1-mm inner diameter, 150-mm length) at a flow rate of 0.45[mL[min^−1^ with a gradient from 100% solvent A (0.1% ammonium formate) to 100% solvent B (methanol). Data are representative of two independent biological replicates.

### Chemical synthesis of p3diT and related nucleotide signal standards

Chemical synthesis of p3diT, sodium salt, and p2diT, sodium salt, were performed as follows: 150 µmol of 5′-Phosphothymidylyl-(3′ −> 5′)-thymidine (p1diT aka pTpT / 5′-pTpT), triethylammonium salt (Biolog Life Science Institute GmbH & Co. KG, cat no. P 129) were suspended in 1,500 µL dried dimethylformamide (DMF), 8 eq. carbonyldiimidazole (CDI) were added and the resulting suspension was stirred under argon for 90 min until all starting material was consumed (analyzed by analytical HPLC). Afterwards, 10 eq. Bis(tri-n-butylammonium) pyrophosphate dissolved in 1500 µL dried DMF were added under argon and stirring was continued for 33 hours at ambient temperature. After this period, the reaction mixture was cooled in an ice batch and was hydrolyzed by the addition of 6 mL water and 6 mL triethylamine (TEA) under stirring. After 10 minutes, the ice bath was removed and stirring continued for a further 2.5 hours at ambient temperature. Another portion of 2 ml each of TEA and water was then added. After 3 hours the hydrolyzed reaction mixture was diluted with 25 mL water and was extracted 3 times with 25 mL methyl tert-butyl ether (MTBE). The product-containing aqueous phase was concentrated by evaporation under reduced pressure to remove traces of MTBE and stored frozen at −20°C until further operations.

The aqueous phase was diluted to 900 mL with water, filtered with a 0.45 µm regenerated cellulose (RC) filter and applied to a Q Sepharose Fast Flow anion exchange column (40–165 µm; 100 × 35 mm) Cl-form, previously regenerated with 2 M sodium chloride and washed with water. The column was washed with water (250 mL), followed by a gradient of 0–0.5 M sodium chloride (NaCl, pH 7, 4,000 mL) in water (detection wavelength 254 nm). The title compounds eluted in separate fraction pools between 800–2,000 mL of the gradient. p3diT-containing fractions were carefully concentrated to a final volume of approximately 30 mL with a rotary evaporator equipped with a drop catcher in-vacuo. Subsequent purification and desalting of p3diT was accomplished by preparative reversed phase medium pressure liquid chromatography (MPLC). The product solution was applied to a YMC Triart Prep C18-S, 12 nm, S-15 µm pre-column (15 µm; 90 × 25 mm) connected to a YMC Triart Prep C18-S, 12 nm, S-15 µm column (15 µm; 480 × 25 mm), previously washed with ethanol and equilibrated with water. Elution was performed with water.

To generate the sodium salt form of p3diT, pooled product-containing fractions were partially concentrated under reduced pressure and loaded onto a Toyopearl SP-650M cation exchange column (65 µm; 110 × 25 mm) Na^+^-form, previously regenerated with 2 M sodium chloride and equilibrated with water. The column was operated with water for elution until no more UV absorption was detectable at 254 nm. After filtration and careful evaporation under reduced pressure, 32.7 µmol p3diT, sodium salt, were isolated with a purity of 99.89% HPLC (theoretical yield: 21.8%).

p3diT:

Formula (free acid): C_20_H_30_N_4_O_21_P_4_ (MW 786.36 g mol^−1^)

UV-Vis (water pH 7.0): λ_max_ 267 nm; ε 17300.

ESI-MS pos. mode: m/z 787 (M+H)^+^, m/z 809 (M+Na)^+^.

ESI-MS neg. mode: m/z 785 (M-H)^−^, m/z 807 (M-2H+Na)^−^.

The p2diT-containing fractions from anion-exchange chromatography were purified and desalted, then transferred to the sodium salt, essentially as described above for p3diT. Following this procedure, 63 µmol of p2diT sodium salt were isolated with 99.48% HPLC purity (theoretical yield: 42%).

p2diT:

Formula (free acid): C_20_H_29_N_4_O_18_P_3_ (MW 706.39 g mol^−1^)

UV-Vis (water pH 7.0): λ_max_ 267 nm; ε 17300.

ESI-MS pos. mode: m/z 707 (M+H)^+^, m/z 729 (M+Na)^+^.

ESI-MS neg. mode: m/z 705 (M-H)^−^.

## Statistics and reproducibility

Experimental details regarding replicates and sample size are described in the figure legends.

## Acknowledgements

The authors are grateful to members of the Kranzusch laboratory for helpful comments and discussion. The work was funded by grants to P.J.K. from the Pew Biomedical Scholars program, the Burroughs Wellcome Fund PATH program, The G. Harold and Leila Y. Mathers Charitable Foundation, The Mark Foundation for Cancer Research, the Cancer Research Institute, the Parker Institute for Cancer Immunotherapy, the Massachusetts Consortium on Pathogen Readiness (MassCPR), and the National Institutes of Health (1DP2GM146250-01), S.Y. is supported by a JSPS Overseas Research Fellowships (202360072) and a Human Frontiers Science Program Long-Term Fellowship (LT0051). D.R.W. is supported through a Helen Hay Whitney Foundation postdoctoral fellowship. X-ray data were collected through support by an agreement between the Advanced Photon Source, a U.S. Department of Energy (DOE) Office of Science user facility operated for the DOE Office of Science by Argonne National Laboratory under Contract No. DE-AC02-06CH11357, and the Diamond Light Source, the U.K.’s national synchrotron science facility, located at the Harwell Science and Innovation Campus in Oxfordshire. X-ray data were additionally collected at The Center for Bio-Molecular Structure (CBMS) that is primarily supported by the NIH-NIGMS through a Center Core P30 Grant (P30GM133893), and by the DOE Office of Biological and Environmental Research (KP1607011). NSLS2 is a U.S. DOE Office of Science User Facility operated under Contract No. DE-SC0012704. This publication resulted from the data collected using the beamtime obtained through NECAT BAG proposal # 311950. Cryo-EM data were collected at the Harvard Cryo-EM Center for Structural Biology at Harvard Medical School.

## Author Contributions

The study was designed and conceived by S.Y. and P.J.K. Phage defense, biochemical experiments, crystallography and cryo-EM structural biology experiments and modeling were performed by S.Y. Phage escape mutant analysis was performed by S.Y. and S.G.F. Nucleotide product LC-MS analysis was performed by S.Y. with assistance from D.R.W. Synthetic nucleotide product synthesis and characterization experiments were performed by M.L. and F.S. The manuscript was written by S.Y. and P.J.K. All authors contributed to editing the manuscript and support the conclusions.

## Competing Interests

M.L. and F.S. are employed at Biolog Life Science Institute GmbH & Co. KG which may sell p3diT and related compounds as research tools. The other authors declare no competing interests.

## Additional Information

Correspondence and requests for materials should be addressed to P.J.K. All illustrations were created using Adobe Illustrator.

## Data Availability Statement

Coordinates and density maps of the *Salmonella* CloA apo, CloA co-expressed with CloB, CloA^HEAA^–dGTP–p3diT, and CloA^HEAA^–dGTP–dTTP complex have been deposited in PDB and EMDB under the accession codes 8P9S, 8P9T, 8P9U, and 8P9V, and EMD-71386, EMD-71388, EMD-71389, and EMD-71390. Coordinates and structure factors of the *Shewanella putrefaciens* dGTPase have been deposited in PDB under the accession codes 9P8W.

## Extended Data Figure Legends

**Extended Data Figure 1 | Discovery of Clover anti-phage defense system.**

**a**, Phylogenetic analysis of ∼650 CloB homologs identified in the IMG database. The inner segments are colored according to identity of the operon encoded adjacent effector protein and the surrounding rings depict the genera of bacteria encoding CloB. **b**, Heatmap illustrating fold defense of *E. coli* expressing Clover system from *Salmonella enterica* and *Escherichia coli* H5. Reduction in plaque forming units (PFU) of wild type CloA or CloA H116A/H117A mutants compared to bacteria expressing a GFP control vector (n=2). **c**, Representative plaque assays of Clover, CloA catalytic mutant, or CloB catalytic mutant operons from *E. coli* cells expressing Clover from *Salmonella enterica* SA20044414 or *Escherichia coli* H5. **d**, Cartoon representation of *Salmonella* CloA octameric assembly formation (top). Overall structure of the octameric CloA assembly (bottom, left) and an example CloA tetrameric unit showing the interface between two dimeric units (bottom, right). **e**, **f**, and **g**, Overview of the CloA_A_-CloA_B_, CloA_A_-CloA_c_, CloA_A_-CloA_D_ interfaces that mediate octameric assembly and detailed view of interacting residues. **h**, Cartoon representation of *Shewanella putrefaciens* CN-32 hexameric assembly (top). Overall structure of the hexameric *Sp*dGTPase assembly (bottom, left) and an example *Sp*dGTPase tetrameric unit showing the interface between two dimeric units (bottom, right). **i**, Cryo-EM structure of a canonical dGTPase *E. coli* Dgt (PDB id: 6OIY) and view of Dgt dimer.

**Extended Data Figure 2 | Cryo-EM data processing for CloA apo structure.**

**a**, *Se*CloA cryo-EM particle picking and classification strategy. **b**, Example motion-corrected micrograph, subjected to particle picking and further analysis. **c**, Example 2D class averages from the particle curation stage. **d**, Gold-standard Fourier shell correlation (GSFSC) curves after FSC-mask auto-tightening, as produced by CryoSPARC. **e**, Local resolution of the final *Se*CloA map.

**Extended Data Figure 3 | CloA activity is stimulated by elevated dNTP levels.**

**a**, AlphaFold3 modelled structures of phage Bas28 dNMP monophosphatase (left) and Bas28 dNMP monophosphatase escape mutant with an S201 nonsense mutation (right). **b**, Cryo-EM structure and magnified view of the active site of the *E. coli* ribonucleotide diphosphate reductase subunit A (NrdA) dimer (PDB id: 6W4X) compared to an AlphaFold3 modelled structure and magnified view of the active site of phage T5 NrdA (right). **c** and **d**, Changes in intracellular dGTP and dTTP concentrations in *E. coli* cells expressing *Salmonella enterica* Clover wild-type or CloA catalytic operon mutant (H117A/D118A) (CloA^HDAA^) operons and infected with phage T5^WT^ or the escape mutant phage T5^NrdA G330^. **e**, LC-MS analysis of purified *Escherichia coli* H5 CloA incubated with dNTPs demonstrates that CloA is a dGTPase activated by the presence of dTTP.

**Extended Data Figure 4 | The nucleotide signal p3diT restrains Clover defense.**

**a**, *Se*CloA–p3diT complex particle picking and classification strategy. **b**, Gold-standard Fourier shell correlation (GSFSC) curves after FSC-mask auto-tightening, as produced by CryoSPARC. **c**, Local resolution of the final CloA–p3diT map. **d**, A_260_ chromatograph of the nucleotide signal released upon heat denaturation of CloA purified from cells co-expressing CloB or CloB^DDAA^ (left). The purified CloB nucleotide signal was further treated with apryrase or a combination of phosphatase (CIP) and nuclease P1. **e**, **f**, and **g**, LC-MS analysis run in negative mode of the purified CloB nucleotide signal alone or treated with apyrase or a combination of CIP and nuclease P1. Formate and chloride ions formed the major adducts [dT+formate]^−^ and [dT+Cl]^−^ respectively observed with deoxythymidine. **g**, Left, comparison between chemically synthesized p3diT, p2diT, p1diT, and the purified CloB nucleotide immune signal. Right, LC-MS analysis run in negative mode of chemically synthesized p3diT, p2diT, and p1diT mixed with purified CloB nucleotide signal. **h**, A_260_ chromatograph of the nucleotide signal released upon heat denaturation of CloA purified from cells co-expressing *Salmonella* or *Xanthomonas* CloB.

**Extended Data Figure 5 | Cryo-EM data processing for CloA bound to p3diT or dTTP binding state.**

**a**, *Se*CloA–dGTP–p3diT complex particle picking and classification strategy. **b**, Gold-standard Fourier shell correlation (GSFSC) curves after FSC-mask auto-tightening, as produced by CryoSPARC. **c**, Local resolution of the final *Se*CloA–dGTP–p3diT complex map. **d**, *Se*CloA– dGTP–dTTP complex particle picking and classification strategy. **e**, Gold-standard Fourier shell correlation (GSFSC) curves after FSC-mask auto-tightening, as produced by CryoSPARC. **f**, Local resolution of the final *Se*CloA–dGTP–dTTP complex map.

**Extended Data Figure 6 | Mechanism of nucleotide signal activation and inhibition.**

**a**, Top, contact helices of p3diT in the *Se*CloA dimer revealing that α2_A_, α4_B_, and α14_A_ helices form the p3diT binding pocket. Bottom, surface carving and electrostatic potential of the p3diT binding pocket. **b**, Top, contact helices of dTTP in the *Se*CloA dimer revealing that α3_A_, α4 _A_, and α18_A_–α21_A_ helices and loop α3_B_–α4_B_ form the dTTP binding pocket. Bottom 180° rotated view of the top figure showing surface carving and electrostatic potential of the dTTP binding pocket. **c**, Top cartoon view of CloA_A_-CloA_B_ dimer in the p3diT bound supressed state structure. Bottom detailed view of α3_A_, α4_A_, and the dGTPase active site in the p3diT bound CloA suppressed state structure. α17_AB_ is omitted for clarity. **d**, Top cartoon view of CloA_A_-CloA_B_ dimer in the dTTP bound active state structure. Bottom detailed view of α3_A_, α4_A_, and the dGTPase active site in the dTTP bound CloA active state structure revealing that dTTP-binding results in a conformational change that stabilizes loop α3_B_–α4_B_ and completes the dGTPase active site. α17_AB_ is omitted for clarity. **e**, Left, superposition of CloA p3diT and dTTP binding states. Middle, 90° rotated view of the left figure of the CloA p3diT binding state. Right, 90° rotated view of the left figure of the CloA dTTP binding state, highlighting additional interactions between loop α3_B_–α4_B_ and Q127_A_ with substrate dGTP. **f**, Rotated view of the superposition of the CloA p3diT and dTTP binding states demonstrates that dTTP binding induces a conformational change that positions the substrate dGTP α-phosphate is proximal to the dGTPase active site residues. **g**, Phosphate release assay measuring dGTP hydrolysis by wild type *Se*CloA and *Se*CloA mutants (Y452A and 3RA) in the presence dTTP and demonstrates that of 200 µM dTTP is sufficient to overcome inhibition induced by 20 µM p3diT. Bar graph depicts the mean, with error bars representing standard deviation.

