## Supplementary figures and images for "Activating and inhibiting nucleotide signals coordinate bacterial anti-phage defense"

### Yamaguchi et al Extended biorxiv (07.09.25).pdf

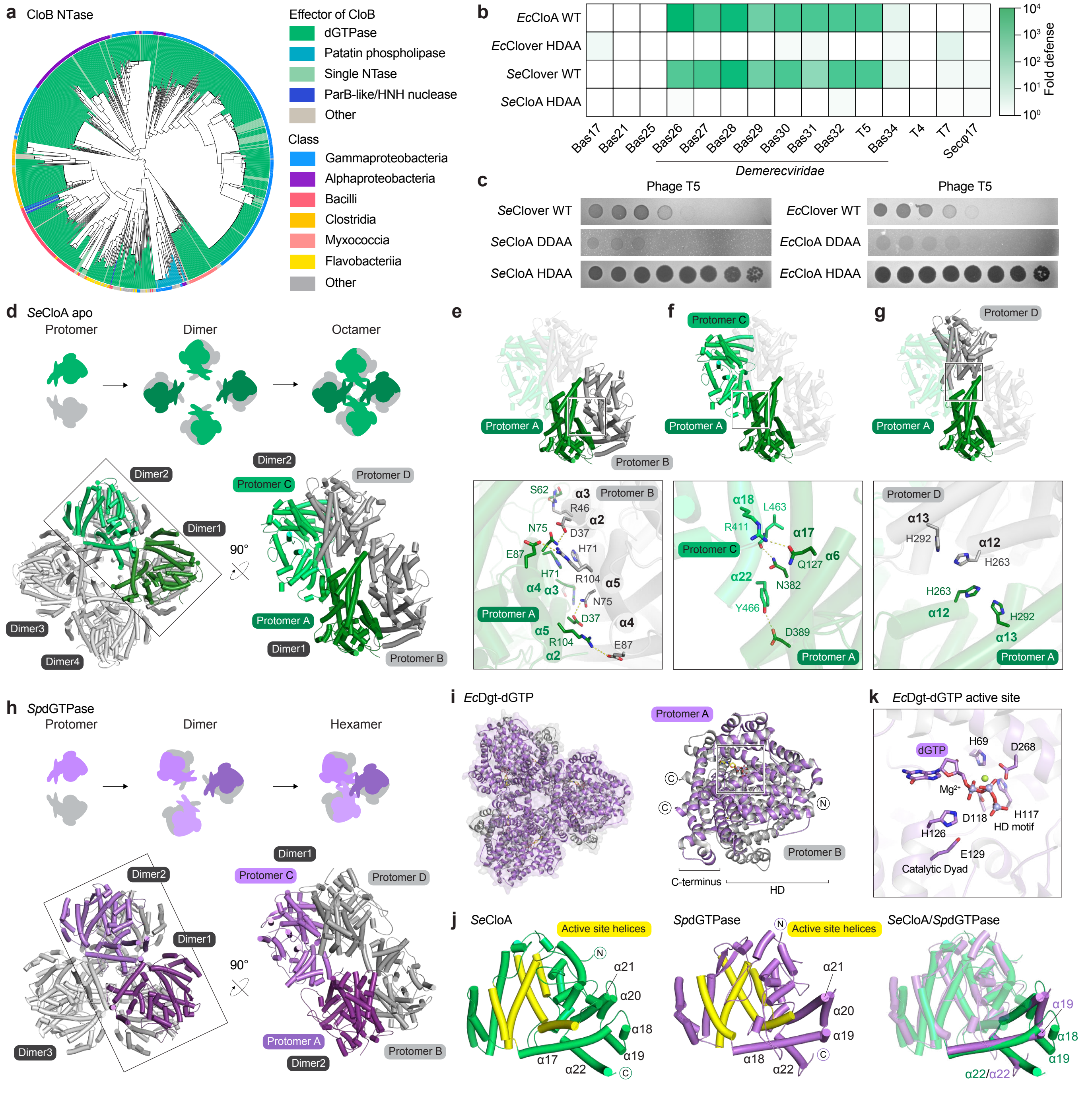

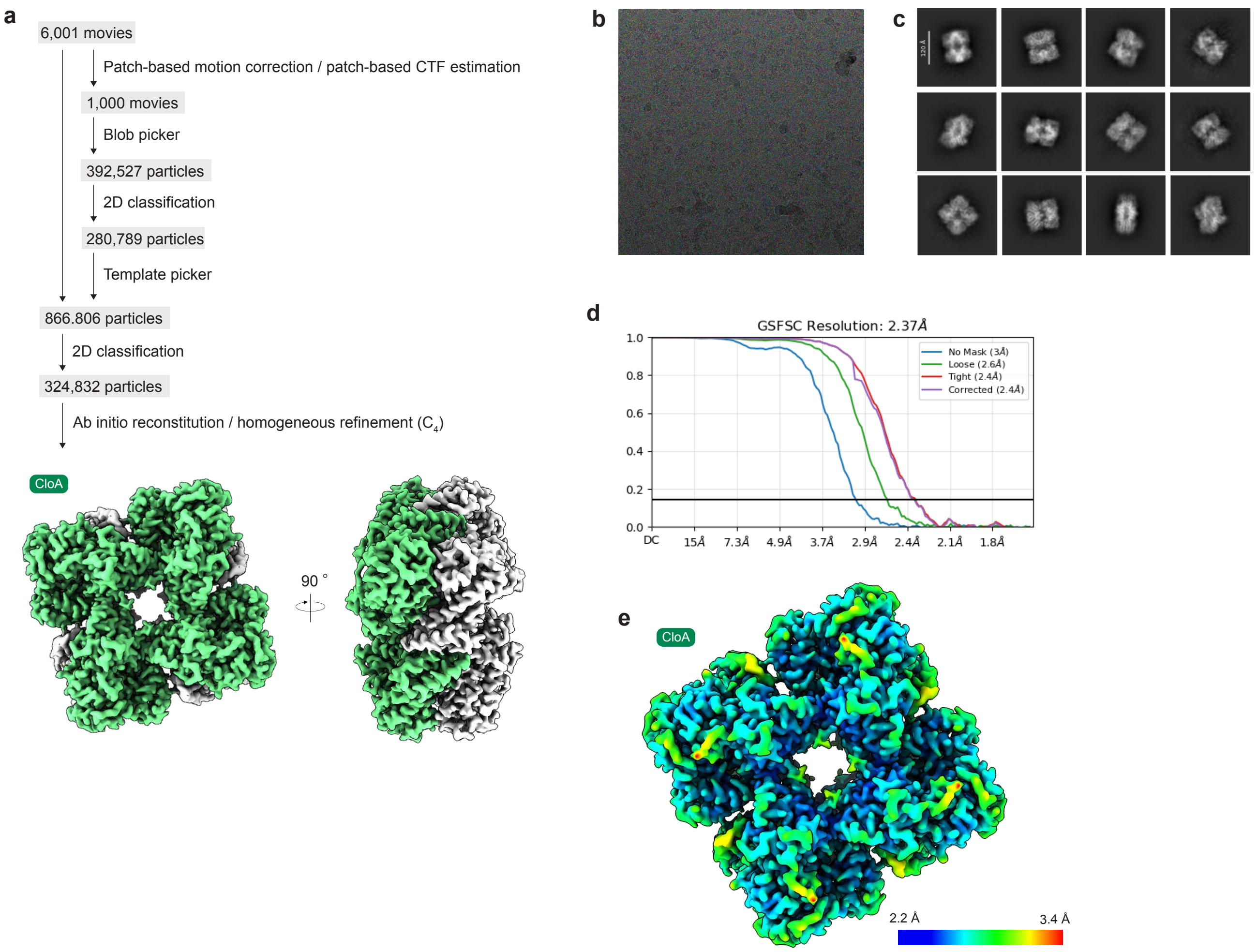

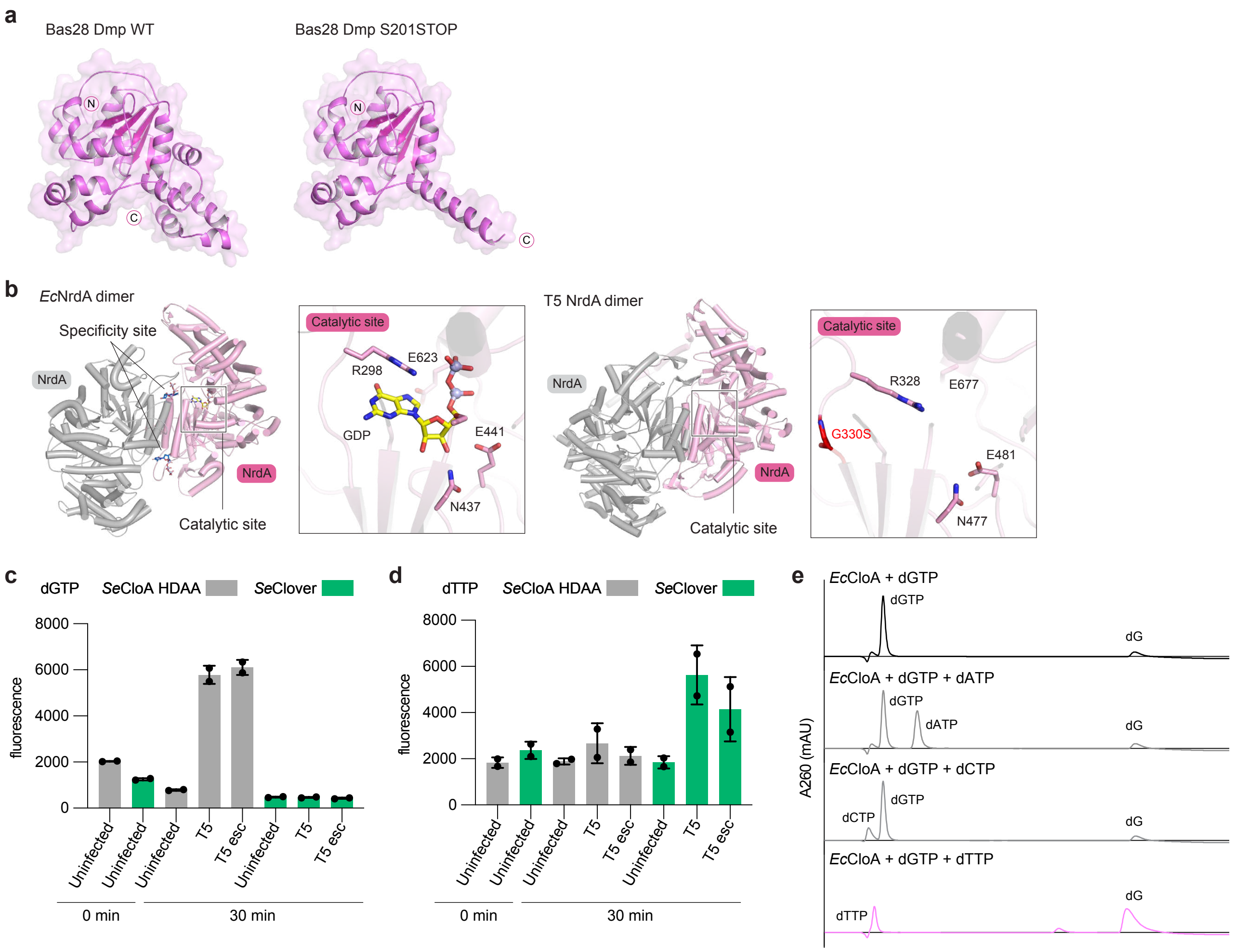

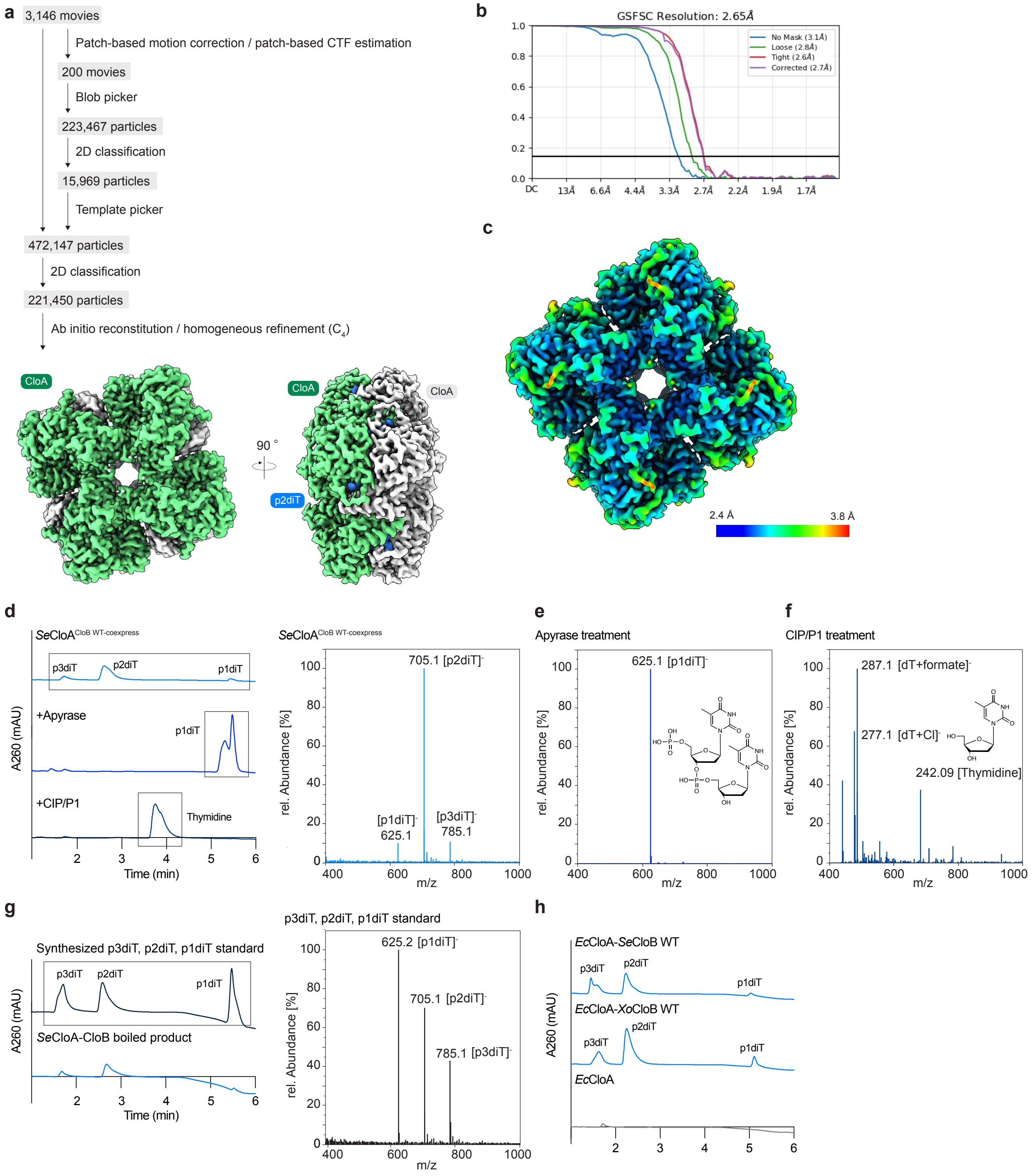

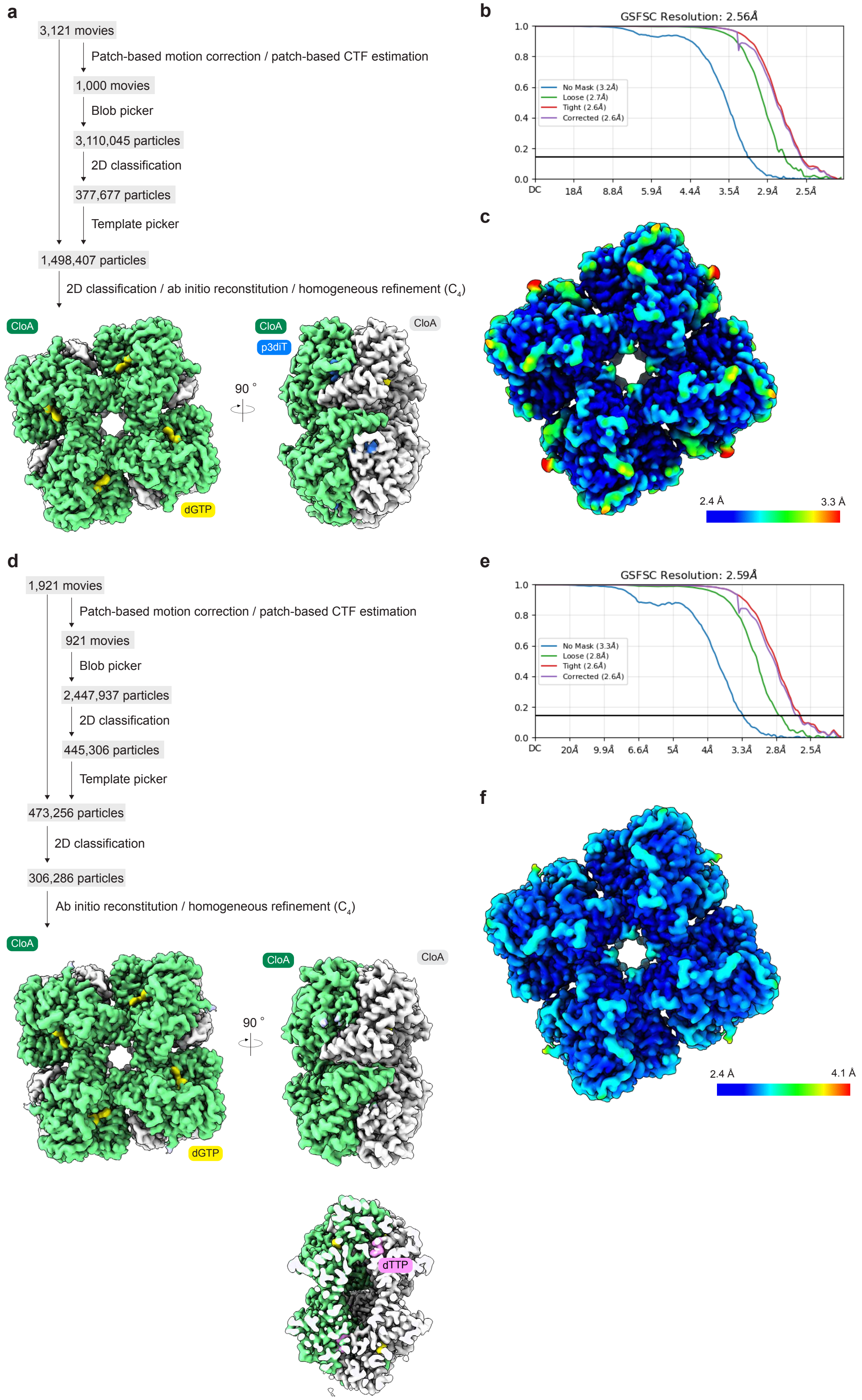

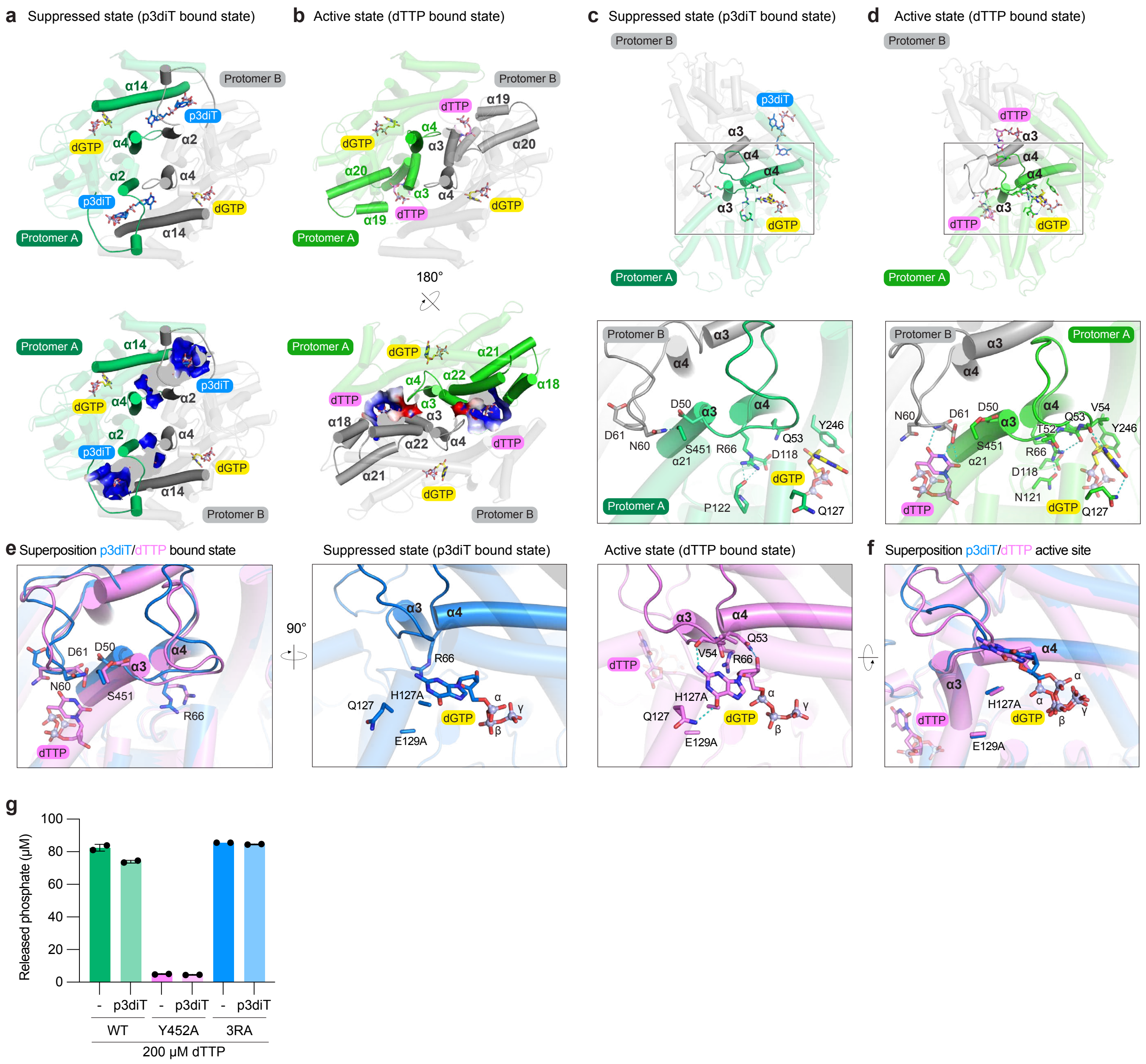
